## Supporting Information for "Sublytic gasdermin-D pores captured in atomistic molecular simulations"

### Supporting Information: Sublytic gasdermin-D pores captured in atomistic molecular simulations

#### TABLE OF CONTENTS


- Supporting Tables S1 - S5
- Supporting Figures S1 - S2
- Supporting Movie Legends S1 - S2

#### Supporting Tables

**Table S1: Asymmetric plasma membrane composition. Fatty acid tails abbreviated as FA.**

| **Lipid** | **FA** | **Full name** | **Charge** | **Inner leaflet [mol %]** | **Outer leaflet [mol %]** |
|---|---|---|---|---|---|
| CCHOL | | Cholesterol | 0 | 40.4 | 40.0 |
| PSM | 18:1/16:0 | N-palmitoyl-D-erythro-sphingosylphosphorylcholine | 0 | 1.0 | 12.0 |
| NSM | 18:1/24:1 | N-nervonoyl-D-oleoyl-sphingosylphosphorylcholine | 0 | - | 9.0 |
| LSM | 18:1/24:0 | N-lignoceroyl-D-oleoyl-sphingosylphosphorylcholine | 0 | - | 8.0 |
| PLPC | 16:0/18:2 | 1-palmitoyl-2-linoleoyl-sn-glycero-3-phosphocholine | 0 | 8.1 | 15.0 |
| SOPC | 18:0/18:1 | 1-stearoyl-2-oleoylphosphocholine | 0 | - | 7.0 |
| PAPC | 16:0/20:4 | 1-palmitoyl-2-arachidonoyl-glycero-3-phosphocholine | 0 | - | 5.0 |
| POPC | 16:0/18:1 | 1-palmitoyl-2-oleoyl-glycero-3-phosphocholine | 0 | 3.0 | - |
| DPPC | 16:0/16:0 | 1,2-dipalmitoyl-glycero-3-phosphocholine | 0 | 2.0 | - |
| PLA20(PE) | 18:0/20:4 | 1-O-stearoyl-2-O-arachidonoyl-glycero-3-phosphoethanolamine | 0 | 11.1 | 3.0 |
| PDoPE | 16:0/22:6 | 1-palmitoyl-2-docosahexaenoyl-glycero-3-phosphoethanolamine | 0 | 8.1 | - |
| SAPE | 18:0/20:4 | 1-stearoyl-2-arachidonoyl-glycero-3-phosphoethanolamine | 0 | 4.0 | - |
| POPE | 16:0/18:1 | 1-palmitoyl-2-oleoyl-glycero-3-phosphoethanolamine | 0 | 3.0 | - |
| PAPS | 16:0/20:4 | 1-palmitoyl-2-arachidonoyl-glycero-3-phosphoserine | -1 | 13.1 | - |
| SAPS | 18:0/20:4 | 1-stearoyl-2-arachidonoyl-glycero-3-phosphoserine | -1 | 1.0 | 1.0 |
| $PI(4,5)P_2$ | 16:0/18:2 | 1-palmitoyl-2-linoleoyl-sn-glycero-3-phosphoinositol-4,5-bisphosphate | -4 | 5.1 | - |

**Table S2: Eisenberg hydrophobicity scores[S1] of the pore facing residues of human GSDMD and mouse GSDMA3 in $\mathrm{kcal\,mol^{-1}}$**

| | human GSDMD | | mouse GSDMA3 | |
|---|---|---|---|---|
| Structural element | Pore facing residues | $\sum$ Eisenberg hydrophobicity | Pore facing residues | $\sum$ Eisenberg hydrophobicity |
| β3 | ADQQSE | -2.73 | MDQQLE | -1.93 |
| β5 | KAGASS | -0.96 | TKKTGS | -2.66 |
| β7 | TKESRS | -4.18 | TNNISP | -1.06 |
| β8 | QEQHSK | -3.76 | LGQSNN | -1.54 |
| $\sum$ full sheet | | -11.63 | | -7.19 |

**Table S3: Restraints used during energy minimization (EM) and stepwise equilibration (steps 1 to 6) of the asymmetric plasma membrane mimetic in $\mathrm{kJ\,mol^{-1}\,nm^{-2}}$.**

| Step | Time [ns] | Timestep [fs] | Ensemble | Lipids | Dihedrals |
|---|---|---|---|---|---|
| EM | | | | 1000 | 1000 |
| 1 | 0.125 | 1 | NVT | 1000 | 1000 |
| 2 | 0.125 | 1 | NVT | 400 | 400 |
| 3 | 0.125 | 1 | NPT | 400 | 200 |
| 4 | 0.5 | 2 | NPT | 200 | 200 |
| 5 | 0.5 | 2 | NPT | 40 | 100 |
| 6 | 0.5 | 2 | NPT | 0 | 0 |

**Table S4: Restraints used during energy minimization (EM) and stepwise equilibration (1 to 3) of the asymmetric plasma membrane mimetic in $\mathrm{kJ\,mol^{-1}\,nm^{-2}}$.**

| Step | Time [ns] | Timestep [fs] | Backbone | Sidechains | Lipid headgroup ($z$) | Water ($z$) |
|---|---|---|---|---|---|---|
| EM | | | 4000 | 2000 | 1000 | 1000 |
| 1 | 5* | 2 | 2000 | 1000 | 10 | 50* |
| 2 | 50 | 2 | 2000 | 1000 | 50 | 0 |
| 3 | 80 | 2 | 1000 | 500 | 0 | 0 |

* For the 27-mer system with an initially lipid filled pore this was extended to 10 ns with a force constant of 1000 $\mathrm{kJ\,mol^{-1}\,nm^{-2}}$ for restraining the z-position of water molecules.

#### Table S5: Atomistic GSDMD$^{NT}$ simulations

| Conformation | Subunits | Simulated time [μs] | Membrane size [$nm^2$] | No. of atoms |
|---|---|---|---|---|
| prepore | 1 | 7.0 | 13.4×13.4 | 222740 |
| prepore | 33 | 3.5 | 38.9×38.9 | 1950552 |
| prepore | 33 | 2.2 | 45.8×45.8 | 3113594 |
| pore | 1 | 5.0 | 12.7×12.7 | 216375 |
| pore | 2 | 5.0 | 12.5×12.5 | 213665 |
| pore | 3 | 5.0 | 12.2×12.2 | 208656 |
| pore | 5 | 4.3 | 18.8×18.8 | 481242 |
| pore | 8 | 2.5 | 25.5×31.9 | 1079514 |
| pore | 10 | 3.0 | 25.4×31.7 | 1071088 |
| pore | 16 | 3.5 | 31.3×31.3 | 1329398 |
| *pore | 16 | 1.4 | 34.9×34.9 | 1497819 |
| pore | 27 | 4.1 | 37.2×37.2 | 2226909 |
| pore | 33 | 5.0 | 37.0×37.0 | 2119785 |
| ◊pore | 33 | 1.5 | 37.0×37.0 | 2119785 |

*simulated in pure DOPC membrane

◊ 70°C continuation of the 37°C simulation after 5μs

#### Supporting Figures

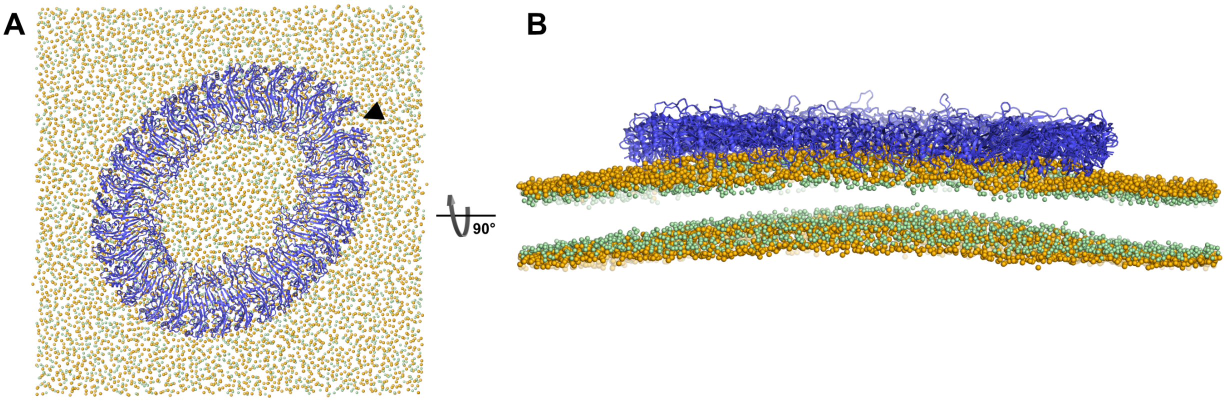


Figure S1: **Snapshot of 33-mer GSDMD$^{NT}$ prepore ring on 46 × 46 nm$^2$ membrane after 2.2 μs of MD simulation, viewed from the top (A) and side (B).** The GSDMD$^{NT}$ backbones are shown in blue cartoon representation. Lipid headgroup phosphates and glycerol oxygen atoms are shown as orange and green spheres, respectively. Water, ions, and lipid tails are not shown for clarity. (A) The arrow indicates where the contacts of the globular domains of two neighboring subunits broke up.

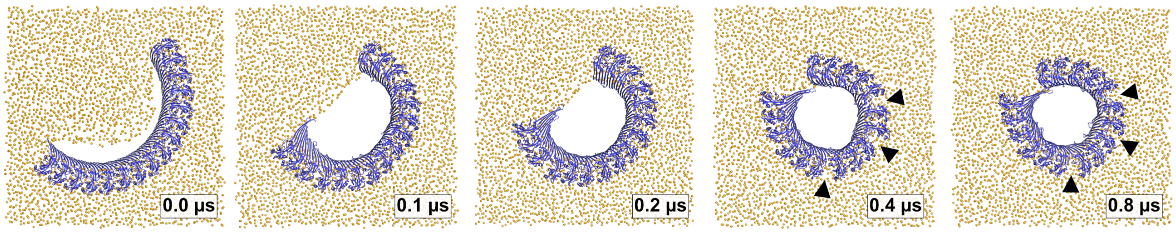


Figure S2: **Time lapse images of pore formation from a 16-meric arc in DOPC membrane.** Top view snapshots of GSDMD$^{NT}$ arcs comprising 16 subunits in pore conformation along MD simulation trajectories show lipids (orange spheres) receding from the inserted β-sheet, before the open protein edges come closer to each other and close a ring shaped pore. The interfaces between globular domains broke in three positions, as indicated with black triangles. Water, ions, and lipid tails are not shown for clarity.

### Supporting Movie Legends

#### Supporting Movie S1

**Formation of a slit-like pore from a membrane inserted 16-meric arc.** Trajectory from 2.0 µs of simulation, showing the plasma membrane edge receding from the protein and then shortening. This draws the ends of the multimer together, causes breakage of the contacts of two neighboring globular domains and ultimately the slit-shaped pore of Figure 4A (2.6 µs) develops. Lipid headgroup phosphates are shown as yellow spheres, cholesterol oxygen atoms as light green spheres, and the protein is shown in blue cartoon representation. Water and ions are not shown for clarity.

#### Supporting Movie S2

**Formation of a ring-like pore from a membrane inserted 27-meric arc.** Trajectory from 4.0 µs of simulation, showing the plasma membrane edge receding from the protein. Lipid efflux first increases the distance between the arc ends, before the open membrane edge then draws the ends together to form the ring-shaped pore of Figure 4B (4.0 µs). Lipid headgroup phosphates are shown as yellow spheres, cholesterol oxygen atoms as light green spheres, and the protein is shown in blue cartoon representation. Water and ions are not shown for clarity.
